# A key role of the EMC complex for mitochondrial respiration and quiescence in fission yeasts

**DOI:** 10.1101/2024.12.16.628658

**Authors:** Modesto Berraquero, Víctor A. Tallada, Juan Jimenez

## Abstract

In eukaryotes, oxygen consumption is mainly driven by the respiratory activity of mitochondria, which generates most of the cellular energy that sustains life. This parameter provides direct information about mitochondrial activity of all aerobic biological systems. Using the Seahorse analyzer instrument, we show here that deletion of the *oca3/emc2* gene (*oca3Δ*) encoding the Emc2 subunit of the ER membrane complex (EMC), a conserved chaperone/insertase which aids membrane protein biogenesis in the ER, severely affects oxygen consumption rates and quiescence survival in *Schizosaccharomyces pombe* yeast cells. Remarkably, the respiratory defects of the *oca3Δ* mutation (EMC dysfunction) is rescued synergistically by the disruption of ergosterol biosynthesis (*erg5Δ)* and the action of the membrane fluidizing agent tween 20, suggesting a direct role of membrane fluidity and sterols composition in mitochondrial respiration in the fission yeast.

## 1 Introduction

Most of the energy that powers eukaryotic cells is generated by the respiration machinery of the mitochondria, double-membraned cytoplasmic organelles that function as the powerhouse of eukaryotic cells. Along the inner membrane, mitochondria contain all components required for respiration and the production of energy, composed of a series of membrane-bound protein complexes that link electron transfer to proton translocation across the membrane (Moe et al., 2023; Vercellino & Sazanov, 2022). In humans, mitochondrial dysfunctions represent the most common cause of the inherited metabolic disease, a diverse group of inherited dysfunctions that not only affect mitochondrial respiratory chain activity and cellular energy production (Davis et al., 2018; Haque et al., 2024) but also many other processes in which this organelle is involved, such lipid biosynthesis and starvation survival (Takeda et al., 2010; Zuin et al., 2008). It is therefore important to optimize experimental techniques that can reliably quantify mitochondrial activity in human cells, but also in model organisms to better understand the basic biology underlying mitochondrial function. To that end, different oxygen sensing techniques have been developed to measure oxygen consumption in mammalian cells (Hynes et al., 2003), isolated mitochondria (Will et al., 2007) or aerobic bacteria and yeasts (Halasa et al., 2014; Jońca et al., 2021; O’Riordan et al., 2000; Turecka et al., 2023; Wu et al., 2018) and therefore, extrapolate to mitochondrial function.

The fission yeast *Schizosaccharomyces pombe* is an important single-celled organism widely used to study various aspects of eukaryotic cell and molecular biology. This model organism resembles human cells in terms of mitochondrial inheritance, mitochondrial transport, sugar metabolism, mitogenome structure and dependence on mitogenome viability (the petite-negative phenotype), which makes this microorganism an attractive model for mitochondrial research (Dinh & Bonnefoy, 2024; Schafer, 2003). However, common techniques developed for oxygen uptake in this yeast usually involve a relatively larger culture volume (Novak & Mitchison, 1990; Zuin et al., 2008), which makes it difficult to determine the metabolic state of these cells under various conditions. Among the different methods, the Seahorse XF24 analyzer turned out to be one of the most successful instruments for accurate analysis of mitochondrial respiration activity in mammal cells, recently used in budding yeasts too (Kumar et al., In press, 2024). In this study, we applied both the Seahorse XF24 instrument and the Seahorse XF HS mini analyzers, to assess oxygen respiration rates in fission yeast cells.

Mitochondrial structure and function require the proper biogenesis of a number of membrane proteins, which are assisted by the ER membrane complex (EMC) (Guna et al., 2018). This chaperone/membrane insertase can functionally substitute for Oxa1 insertase in mitochondria (Schneider et al., 2022). The structural integrity of EMC depends on the ‘core subunits’ Emc1, Emc2, Emc3, Emc5 and Emc6, which are essential for both assembly and function of the complex (Guna et al., 2018; Shurtleff et al., 2018; Volkmar et al., 2019). In humans, depletion of any of these EMC components leads to mitochondrial dysfunction among other cellular defects (Volkmar & Christianson, 2020). Here, we used the Seahorse XF24 and the Seahorse XF HS mini analyzers, to assess the effects on mitochondrial activity caused by EMC-dysfunction in Oca3/Emc2 -depleted *S. pombe* cells.

## 2 Materials and Methods

### 2.1 Media and growth conditions

Standard fission yeast growth media were employed throughout the experiments as described in (Moreno et al., 1991). Yeast cultures were grown in YES media in 100 ml flasks with shaking at 30ºC (Moreno et al., 1991). Strains used in this study are listed in Table S1.

### 2.2 Fluorescence Microscopy

For live-cell imaging, cells were mounted in an Ibidi μ-Slide 8 Well chamber (Ibidi 80826) adhered to the chamber with soybean lectin (Sigma L1395). Imaging was conducted using a spinning-disk confocal microscope (IX-81, Olympus; CoolSNAP HQ2 camera, Plan Apochromat 100x, 1.4 NA objective, Roper Scientific) and Metamorph software, with the temperature maintained at 30°C. For mitochondrial staining, MitoTracker CMxRos (200 nM in medium from the stock of 1 mM in DMSO, ThermoFisher M7512) was added to exponentially growing *S. pombe* cells and fluoresecence visualized in live cell imaging at the spinning-disk confocal microscope. Maximal projections of 26 slices with a Z-step of 0.3 μm were presented.

### 2.3 Oxygen Consumption using the Seahorse instrument

Plates were prepared the day before the assay with Poly-D-Lysine coating by adding 100 μL of 1:1 diluted Poly-D-Lysine (50 μg/mL) (Sigma P6407) in ultrapure water for 30 minutes at room temperature (RT), then removing the excess and allowing it to dry. Calibration solution was added following Agilent specifications to the plates and incubated at the desired temperature for 24 hours.

On the assay day, cultures were grown to exponential phase (2×10^6^). Carbonyl cyanide 4-(trifluoromethoxy) phenylhydrazone (FCCP,10 mM, Sigma SML2959) and Antimycin A (5 mM, Sigma A8674) stocks were prepared in DMSO and diluted in fresh media: FCCP at 1:1000 (10 μM) and Antimycin A at 1:100 (5 μM).

When using the Seahorse XF24 the plates were prepared as following: The injection plate was loaded with FCCP and Antimycin A volumes: Well A received 56 μL of FCCP (10 μM), Well B received 62 μL of FCCP (10 μM), Well C received 69 μL of Antimycin A (5 μM), and Well D remained empty. For the assay plate, cultures were diluted to a DO600 of 0.1, then 500 μL of fresh medium was added to four blank wells and 500 μL of culture was added to the wells being measured. The plate was centrifuged at 500 g for 3 minutes and then incubated at the desired temperature for 30 minutes to acclimate the cells.

When using the Seahorse XF HS Mini the plates were prepared as following: The injection plate was loaded with FCCP and Antimycin A volumes: Well A received 20 μL of FCCP (10 μM), Well B received 22 μL of FCCP (10 μM), Well C received 24 μL of Antimycin A (5 μM), and Well D remained empty. For the assay plate, cultures were diluted to a DO600 of 0.3, then 180 μL of fresh medium was added to two blank wells and 180 μL of culture was added to the wells being measured. The plate was centrifuged at 500 g for 3 minutes and then incubated at the desired temperature for 30 minutes to acclimate the cells.

Measurements were performed using the corresponding Seahorse instrument during 10 cycles for a 90-minute total. FCCP injections were added in the 4th and 6th cycles and Antimycin A injection at the 8th cycle.

### 2.4 Survival and longevity assays in mitotic and meiotic quiescent cells

We followed the method to determine survival and longevity (Roux et al., 2009). In brief, cells were cultured to late exponential growth phase in rich media (YES) across three independent biological replicates cultures for each mutant strain. We scored the number of cells/ml in liquid media during exponential growth. All strains showed a similar duplication time of 3,5 hours. Cells/mL were accurately counted in a Neubauer chamber under the microscope and proper dilutions were carried out to plate 200 cells on solid YES medium from late exponentially growing cultures to establish the initial cell count survival (point T-1). The remainder of the culture was maintained until the stationary phase was reached, defined as the point at which cell division ceased for two consecutive generation times. Upon achieving stationary phase, 200 cells were plated again (at time point T0), and this procedure was repeated every 24 hours until cells were 0.1% of the original count (T1 to T4). Once the colonies had fully developed, we quantified the number of viable cells.

For spore viability, opposing mating type (h+ or h-) *wild-type* and/or *oca3Δ* cells were mated until zygotes were produced and asci developed. Asci were dissected in YES plates with a Singer micromanipulator MSM400, and viability determined by the formation of colonies from the isolated spores. To follow living sporulation, h+ and h-cells were mated and resulting zygotes were mounted under the fluorescence microscope to follow mitochondria (marked with Arg11-mCherry) and the forespore formation (visualized with Psy-GFP) through meiosis and spore development

## 3 Results and Discussion

### 3.1 Oxygen consumption rates in *S. pombe* cells

Measurements of oxygen consumption rates have been central to the recent resurgent interest in mitochondrial metabolism (Divakaruni & Jastroch, 2022). To this end, the Seahorse analyzer is one of the most successful instruments for the accurate analysis of mitochondrial respiration activity. This instrument was originally designed to monitor oxygen consumption in adherent living cells, but it has been successfully adapted to assay respiration rates in worms (Haroon & Vermulst, 2019) and different yeasts (Vajrala et al., 2021; Zhang et al., 2018). Very recently, it has been proposed as a generalized method for measuring oxygen consumption in Saccharomyces yeasts (Kumar et al., In press, 2024), also applied to *S. pombe* cells (Ohsawa et al., 2024). Therefore, in order to assess mitochondrial activity in diverse genetic backgrounds and chemicals in this model yeast, we decided to adapt the Seahorse bioanalyzer to measure oxygen consumption in *S. pombe* cells.

The analysis can be performed in real time in multi-well plates (8, 24 or 96 wells), so that multiple yeast strains and controls can be analyzed simultaneously. In addition, each well is equipped with ports that can inject different chemical solutions into the wells as needed, so that yeast cells can be exposed to multiple reagents in succession. Accurate measurements of oxygen consumption with this instrument require that the samples are directly beneath the probe. For this purpose, yeast cells are kept attached to the wells of multi-well plates with poly-D-lysine (see methods). To measure oxygen consumption rates we have used both, the Seahorse XF24 for permissive temperature and the Seahorse XF HS mini at low temperatures (20ºC) (see in Fig. 1).

**Figure 1.**
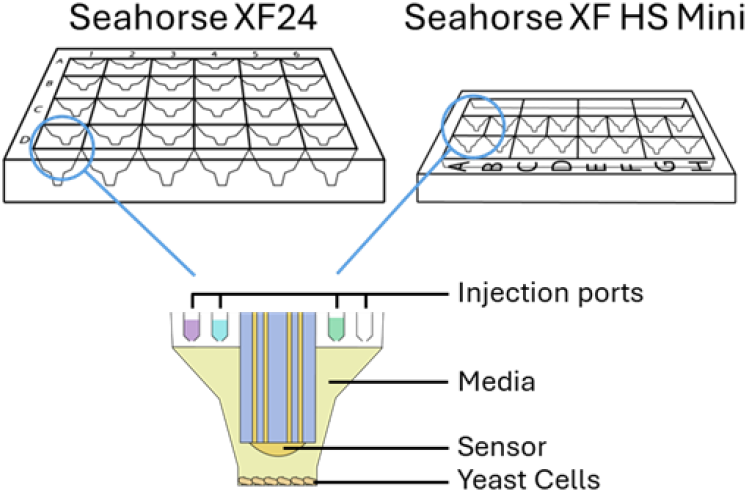
Schematic of the Seahorse XF Bioanalyzer plates. Multi-well cell culture plates housing fission yeast cells (top scheme). Each probe contains injection ports that can be used to add compounds, such as FCCP or antimycin A, and two sensors for oxygen levels and solution pH (bottom scheme).

As shown in Fig. 2, high levels of oxygen consumption are observed in *wild-type S. pombe* cells. Compared to these *wild-type* cells, the *coq4Δ* respiratory-deficient mutant cells used as a negative control show a very poor oxygen consumption rate (Pelosi et al., 2024). Addition of the carbonyl cyanide protonophore 4-(trifluoromethoxy) phenylhydrazone (FCCP) (6 μM) induces maximal respiration in *wild-type* cells by uncoupling electron transport from oxidative phosphorylation (Bertholet et al., 2022; Terada, 1990) with no significant effects in the *coq4Δ* mutant strain (see Figure 2). In contrast, antimycin A is a potent inhibitor of mitochondrial electron transport that can be used to block respiration (Labs et al., 2016). Accordingly, addition of this drug dramatically inhibits oxygen consumption in *wild-type* cells to values close to those of *coq4Δ* respiratory-deficient mutant cells (Figure 2). Similar kinetics are observed after addition of FCCP and antimycin A in both strains (*wild-type* and the *coq4Δ* mutant control) at 20°C and 30°C, but with an oxygen consumption rate about 5-fold lower at the former temperature (Figure 2). These results indicate that oxygen consumption rates measured on the Seahorse analyzer provide accurate estimations of the respiratory activity in *S. pombe* cells.

**Figure 2.**
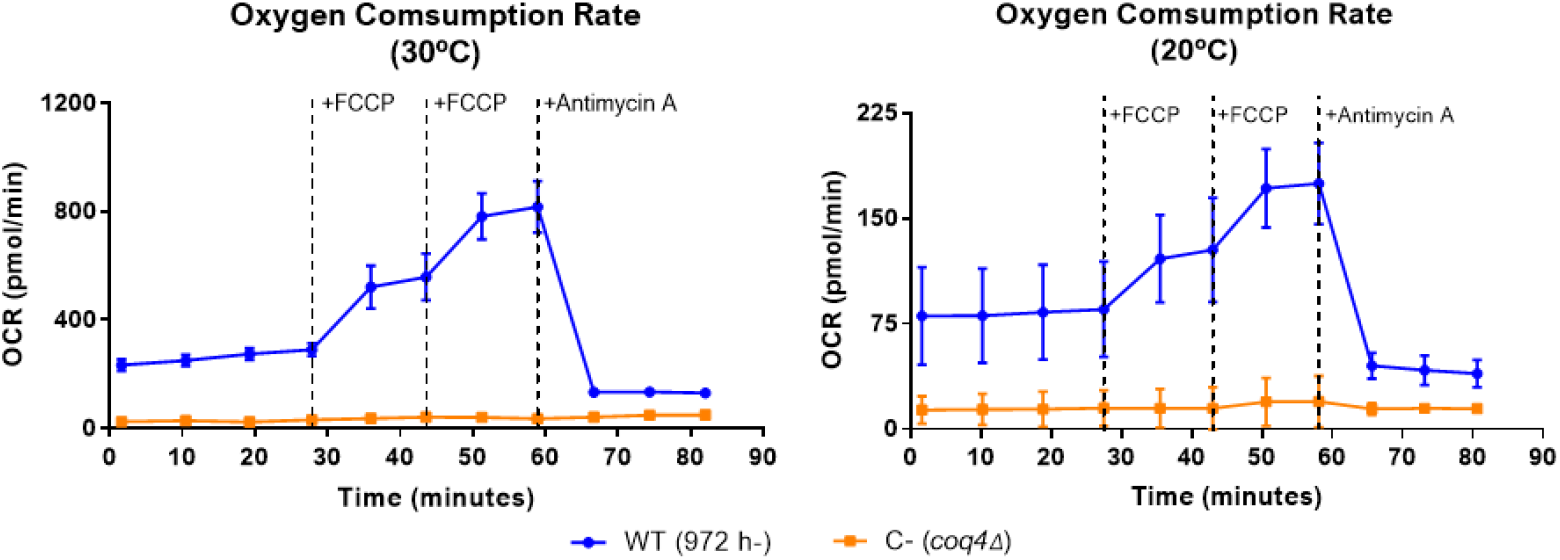
Respiration rates in fission yeast cells in the Seahorse Analyzer. O2 consumption rate (OCR, pmol/min) (Seahorse XF24 measurements at 30ºC and Seahorse XF HS Mini at 20ºC) in *wild-type* (wt) cells at 20ºC and 30ºC growth temperatures. Respiratory-deficient *coq4Δ* mutant cells are used as negative control. Scale at 20ºC is five times lower than 30ºC. FCCP and antimycin A were injected as described in methods.

### 3.2 EMC dysfunction affects the rate of mitochondrial respiration

Biogenesis of membrane proteins in the ER requires the assistance of the highly conserved EMC complex. This ER membrane proteins insertase plays a critical role at ER-mitochondria contact sites in eukaryotic cells (Volkmar & Christianson, 2020). Accordingly, we have recently observed that depletion of Oca3/Emc2 (identified in *S. pombe* cells due to their defects in normal cell cycle proliferation (Tallada et al., 2002), alters mitochondrial structure (Berraquero et al., 2024). Using Mitohealth staining to visualize mitochondrial membranes microscopically, we observed that Oca3/Emc2 depletion affects the normal mitochondrial tubular network of *S. pombe* cells, resulting in aggregates of mitochondrial membranes (see Figure 3A).

**Figure 3.**
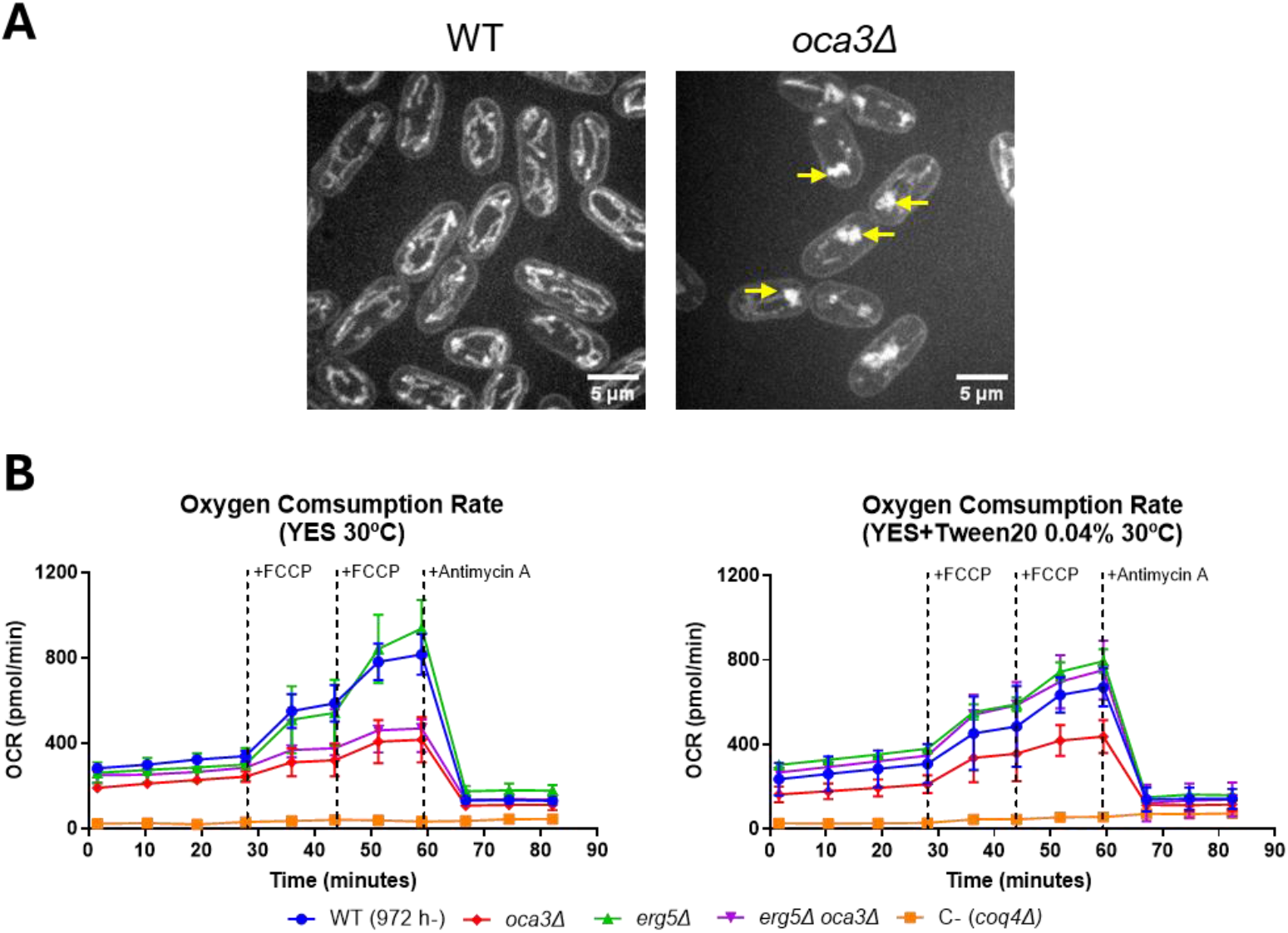
Effects of Oca3/Emc2 depletion on mitochondrial structure and function at 30ºC. A. *in vivo* Mitohealth (ThermoFischer H10295) staining highlights mitochondrial membrane localization in *wild-type* and *oca3Δ* cells. Deletion of *oca3* results in viable mitochondrial aggregation where the mitochondrial network collapses (arrows). **B**. O2 consumption rate (OCR, pmol/min) (Seahorse XF24 measurements) in *oca3Δ, erg5Δ* and the *oca3Δ erg5Δ* double mutant strain at 30ºC (left panel) and the same assay with the addition of tween 20 (0,04%) (right panel). *Wild-type* and respiratory-deficient *coq4Δ* mutant cells are included as positive and negative controls respectively. FCCP and antimycin A were injected as described in methods.

The mitochondrial respiratory chain is a key mechanism of energy production in the overall cellular metabolism of eukaryotic cells, located in the inner mitochondrial membrane (Vercellino & Sazanov, 2022). Therefore, to determine whether the altered mitochondrial architecture of *oca3Δ* cells also leads to mitochondrial respiratory dysfunction, we studied oxygen consumption rates in *S. pombe* cells depleted for Oca3/Emc2. Compared to *wild-type* cells, the oxygen consumption rate is significantly reduced (more than 50% at its maximal level at 30ºC) in the *oca3Δ* strain.

Antimycin A decreases the oxygen consumption rate, but the addition of FCCP does not significantly induce the respiration rate in this strain (Fig. 3B). Therefore, according to these direct indicators of mitochondrial activity, we conclude that the EMC complex is required for full respiratory activity in the fission yeast.

### 3.3 Membrane fluidizing may rescue respiration deficiency of EMC-deficient cells

Mitochondrial membranes are poor in ergosterol content, but this sterol is required at ER-Mitochondrial junctions for proper mitochondrial activity (Volkmar & Christianson, 2020). Disruption of the ergosterol synthesis pathway at the penultimate step in the *erg5Δ* mutant produces ergosta-5-7-dienol instead of ergosterol (Skaggs et al., 1996). As shown in Figure 3B, Erg5-deficient cells respire efficiently, showing oxygen consumption rates like *wild-type* cells. This result suggests that ergosta-5-7-dienol acts on cell membranes in a manner very similar to ergosterol. Cells lacking Oca3/Emc2 over-accumulate ergosterol (Berraquero et al., 2024). Oxygen consumption in the *oca3Δ erg5Δ* double mutant cells is like that of the *oca3Δ* single mutant cells, indicating that the *oca3Δ* mutation is epistatic over *erg5Δ* (Figure 3B), probably by overaccumulation of ergosta-5-7-dienol, the end-product of the ergosterol pathway in the *erg5Δ* strain. Interestingly, addition of tween 20 (0.04%) slightly increases oxygen consumption rates in the *oca3Δ* mutant cells, but addition of this non-ionic detergent fully recovers the cellular respiration capacity to *wild-type* levels in the *oca3Δ erg5Δ* double mutant strain (Figure 3B). Therefore, respiration in *oca3Δ* is rescued synergistically by the disruption of ergosterol biosynthesis (*erg5Δ*) and the membrane fluidizing agent tween 20 (see in Figure 3), perhaps by recovering the optimal membrane fluidity necessary for mitochondrial activity (Berraquero et al., 2024).

### 3.4 EMC-deficiency disturbs entry into quiescence in *S. pombe* cells

Survival of cells facing limited resources and other stressing environments depends on mechanisms regulating entry into a quiescent state, a temporary and reversible exit from proliferative growth. Proper mitochondrial function has been shown to be essential in this process (Takeda et al., 2010; Zuin et al., 2008). Given the severe defects in mitochondrial structure and respiration observed in the Oca3/Emc2-depleted mutant, we assessed survival of *oca3Δ* cells following entry into the G0 phase of the cell cycle, a quiescent state which allows cells to survive during months, and in the formation of spores through meiosis, a process of the life cycle that maintains survival of yeasts for years (Sun & Gresham, 2021).

To analyse the effects of EMC-dysfunction in the transition to G0 phase, we followed the method described in Roux et al., 2009. We grew *wild-type* and *oca3Δ* cells in YES medium at 30ºC until stationary phase and the number of viable cells (colony forming units out of 200 plated cells) was determined in exponentially growing cells (T-1, Figure 4A left), at early G0 (T0, second generation time with no cell number increase) and every 24 hours (see materials and methods). Despite reaching the stationary phase at similar values, the viability of *oca3Δ* mutant cells fell down much faster than the *wild-type* as cells entry into quiescence (T0, Figure 4A left). However, at longer periods (up to four days in our study -T1 to T4), survival of *wild-type* and *oca3Δ* cells exhibit similar values, suggesting that the chronological lifespan of cells that survive entry into G0 is similar in both strains (Figure 4A right). These results emphasize the importance of the EMC complex in mitochondrial function, which thereby influences the transition between proliferative and quiescent states in the vegetative life cycle of *S. pombe* cells.

**Figure 4.**
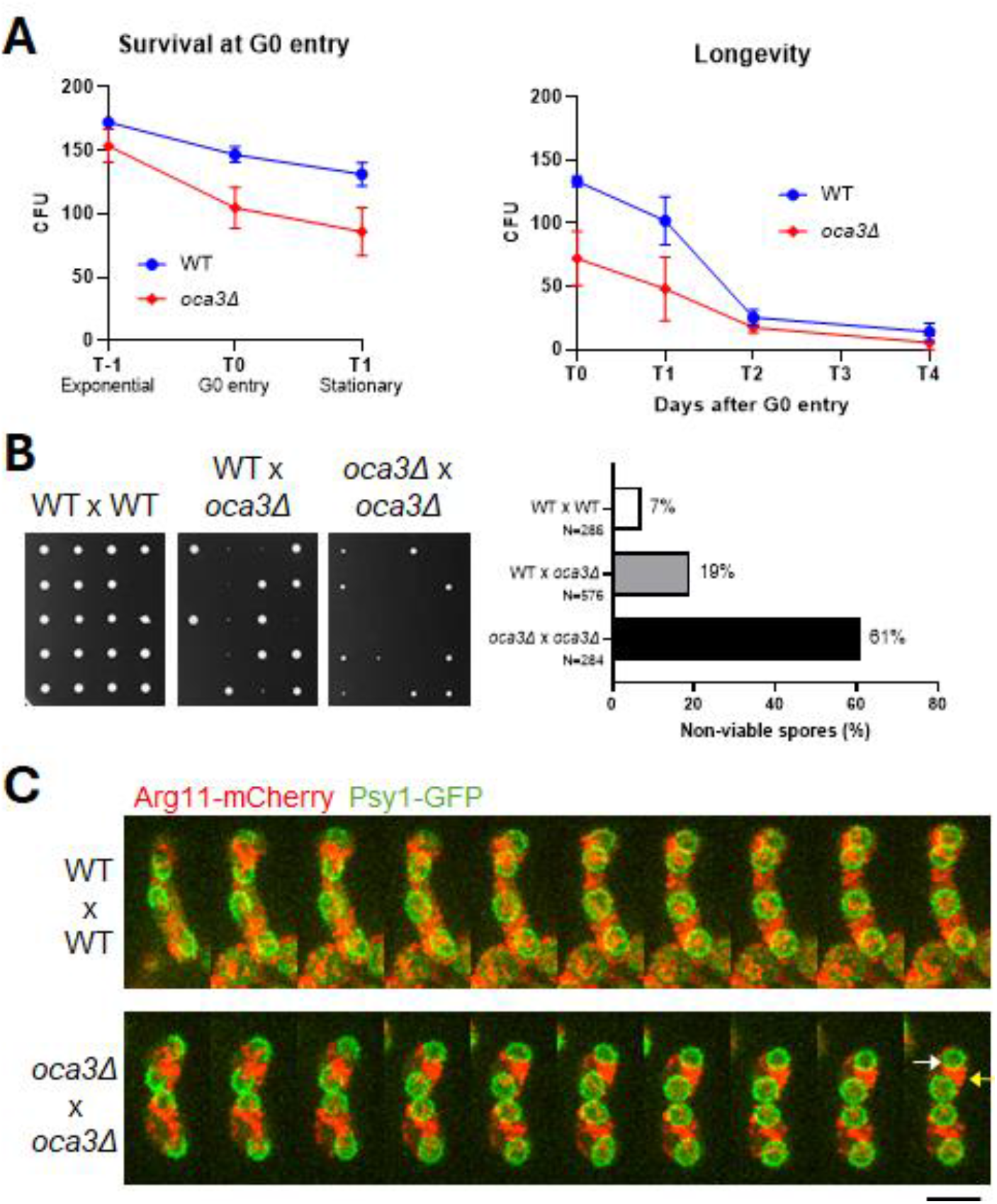
Effects of mitochondrial dysfunction caused by EMC-deficiency on mitotic and meiotic quiescence cells. A. Entry into the stationary phase (T-1: late exponential growing cells, T0: cells entering stationary phase: T1: G0 after 24h). Chronological lifespan assays of *wild-type* and *oca3Δ* cells (right panel). **B**. Isolated spores from dissected asci (tetrads) of zygotes from the indicated mating cells (left panel). Mitochondrial dysfunction yields smaller colonies in EMC-deficient spores (2:2 segregation in zygotes from WT x *oca3Δ*). Lethality (%) of these spores is indicated (right panel). Spores were germinated in YES agar plates until colonies were observed. **C**. Time-lapse images of representative sporulating zygotes from the mating of *wild-type* (WT x WT) (upper panel) and *oca3Δ x oca3Δ* (lower panel) cells. Forespore formation (Psy1-GFP) and mitochondria (Arg11-mCherry) are visualised simultaneously. Mutant meiosis give rise to spore with no visible mitochondria (white arrow) and aggregated mitochondria outside the spores (yellow arrow). Each image is acquired every 8 minutes from a maximal projection of 26 slices of 0.3 μm. Scale bar 5 μm.

The spores of yeasts are quiescent cells that are highly resistant to various stresses, such as heat, digestive enzymes, and organic solvents (Egel, 1977). To determine how mitochondrial dysfunction of EMC-deficient cells influenced spore viavility, we determined the viability of spores isolated from disected tetrads in asci originated when mating: i. *wild-type* x *wild-type* cells (control), ii. *wild-type* x *oca3Δ* cells, and iii. *oca3Δ* x *oca3Δ* cells. As shown in Figure 4B, the fraction of spores that germinate to form a colony is highly reduced in the meiosis of heterozygous zygotes, and severelly affected in meiosis from *oca3Δ* x *oca3Δ* homozygous zygotes.

We have recently shown that segregation/stability of mitochondrial DNA is affected by the mitochondrial aggregates formed in EMC-deficient cells (Berraquero et al., 2024). To determine whether this struture disturbs mitochondrial segregation in meiosis, live imaging of sporulating zygotes with labeled mitochondria (Arg11-mCherry) and a forespore marker (Psy-GFP) were visualized under the microscope during meiosis. As expected, all mitochondria are efficiently distributed among the four meiotic products in zygotes from *wild-type* mating cells (N=53). However, this organele missegreates when mating *oca3Δ* cells. In *oca3Δ* x *oca3Δ* zygotes, most of the mitochondrial mass remaining outside of the spore during forespore development, leading to 18% of spores without visible mitochondria (N=100) (Figure 4C, Video S1 and S2). *S. pombe* cells are petite-negative yeasts, and under *wild-type* genetic backgrounds mitochondrial DNA and functions are required for survival (Schafer, 2003). Therefore, the lethality of meiotic products in zygotes harbouring the *oca3Δ* mutation are likely due to the severe deffects in mitochondrial segregation and/or the content/quality of the mitochondria inherited during meiosis of EMC-defficient cells.

Overall, we show here that measurement of oxygen consumption with the Seahorse XP Analyzer can help to identify mitochondrial mechanisms of action upon pharmacological and genetic interventions to accurately assess mitochondrial function. Our results indicate that the EMC complex, a conserved ER chaperone/insertase that assist biogenesis of membrane proteins at ER-Mitochondrial contacts (Volkmar & Christianson, 2020) is required for full mitochondrial respiration activity, as well as to attain proper mitochondrial activity and structure to permit the transition to a quiescence state in mitotic and meiotic cells.

## Acknowledgments

We thank Katherina García for her assistance in the advanced microscopy facility, Victor Carranco for excellent technical assistance, Carlos Santos for his suggestions on the Seahorse XF24 Analyzer protocols, and all members of the yeast genetics group at the CABD for valuable comments and discussions. This work was supported by the Spanish Ministerio de Ciencia e Innovación (grant numbers PID2019-111124GB-I00 to J.J.).

## Authors contributions

M.B., V.A.T. and J.J. designed the experiments. M.B. performed the experiments and analyzed the data. V.A.T. analyzed the data and supervised experimental protocols. J.J. analyzed the data, acquired funding and wrote the paper with input from M.B. and V.A.T.

## Supporting information

**Table S1.** List of strains used in this study.

| Strain | Source | Genotype |
| --- | --- | --- |
| VA1 | VAT Collection | h- 972 |
| VA240 | VAT Collection | h+ his- |
| JJ2374 | JJ Collection | h+ oca3::Kan |
| JJ2375 | JJ Collection | h- oca3::Kan |
| JJ2426 | FY33691 NBRP, Japan | h+ coq4::kanMX6 leu1-32 ura4-D18 |
| JJ2493 | FY17341 NBRP, Japan | h- erg5::ura ura- leu- |
| JJ2508 | JJ Collection | h? erg5::ura oca3::kan |
| JJ2606 | Daga R. Lab | h+ arg11-mCherry:nat |
| JJ2607 | JJ Collection | h- oca3::kanMX4 arg11-mCherry:nat |
| JJ2785 | Daga R.. Lab | h90 GFP-psy1:leu leu- |
| JJ2786 | This study | h+ GFP-psy1:leu leu- |
| JJ2787 | This study | h- GFP-psy1:leu leu- |
| JJ2790 | This study | h+ GFP-psy1:leu oca3::kanMX4 arg11-mcherry:nat |
| JJ2791 | This study | h- GFP-psy1:leu oca3::kanMX4 arg11-mcherry:nat |
| JJ2793 | This study | h+ GFP-psy1:leu arg11-mcherry:nat |
| JJ2794 | This study | h- GFP-psy1:leu arg11-mcherry:nat |

Video S1. Time-lapse of sporulating zygotes (representative ascus) from the mating of *wild-type* (WT) (left video) and *oca3Δ* (right video) cells. Each frame is acquired every 4 minutes. Brightfield shows cells and mitochondria in red (Arg11-mCherry) channels from max projection of 26 slices of 0.3 μm.

Video S2. Time-lapse of sporulating zygotes (representative ascus) from the mating of *wild-type* (WT) (left video) and *oca3Δ* (right video) cells. Each image is acquired every 4 minutes from a max projection of 26 slices of 0.3 μm. Forespores (Psy1-GFP) are green and mitochondria (Arg11-mCherry) red.

