## Supplementary figures and images for "A key role of the EMC complex for mitochondrial respiration and quiescence in fission yeasts"

### Supplemental Figs S1

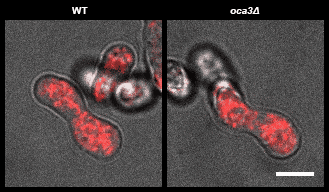
